# Species-specific satellite DNA composition dictates PRC1-mediated pericentric heterochromatin

**DOI:** 10.1101/2024.10.11.617947

**Authors:** Piero Lamelza, Malena Parrado, Michael A. Lampson

## Abstract

Pericentromeres are heterochromatic regions adjacent to centromeres that ensure accurate chromosome segregation. Despite their conserved function, they are composed of rapidly evolving A/T-rich satellite DNA. This paradoxical observation is partially resolved by epigenetic mechanisms that maintain H3K9me3-based heterochromatin, independent of specific DNA sequences. However, these mechanisms can function only after H3K9me3 has already been established, and this mark is absent from paternal chromatin in the mouse zygote. It is unknown how variation in satellite DNA sequence impacts alternative forms of heterochromatin at this earliest stage of life. Here we show functional consequences of satellite DNA variation for pericentric heterochromatin formation, recruitment of the Chromosome Passenger Complex (CPC), and interactions with the mitotic spindle. The AT-hook of Polycomb Repressive Complex 1 (PRC1) directly recognizes A/T-rich satellite DNA and packages it in H2AK119ub1 heterochromatin. By fertilizing *M. musculus* eggs with sperm from other mouse species, we show that divergent satellite sequences differ in their ability to bind PRC1, resulting in differences in H2AK119ub1 heterochromatin formation on mitotic chromosomes. Furthermore, we find that satellites that robustly form H2AK119ub1 inhibit molecular pathways that recruit the CPC to pericentromeres, increasing microtubule forces on kinetochores during mitosis. Our results provide a direct link between satellite DNA composition and pericentromere function and highlight early embryogenesis as a critical point in development that is sensitive to satellite DNA evolution.

## Introduction

Satellite DNA is composed of tandemly repeated DNA sequences that make up large non-coding regions of eukaryotic genomes, e.g. 7% and 8% of human and mouse genomes, respectively ^1,2^. Satellites are among the most rapidly evolving genomic sequences, with their monomer size, nucleotide sequence, genomic distribution and abundance varying widely between species^3,4^. Despite these rapid changes, they often underlie chromosomal loci with highly conserved functions^5,6^. Recent advances in long-read sequencing technology have made huge progress in precisely characterizing the genetic composition and diversity of repetitive satellite arrays^1,7–9^. However, it is still largely unknown what functional consequences may arise from the natural diversity of these sequences.

Centromeres and pericentromeres, two loci with conserved functions in orchestrating accurate chromosome segregation during cell division, are composed of rapidly evolving satellite arrays in many species. This paradoxical observation is partially resolved by evidence indicating that centromere and pericentromere function, packaging and inheritance rely on epigenetic mechanisms. The histone H3 variant CENP-A (or CenH3) defines centromeres in most eukaryotes^10^, while pericentric satellites are enriched with H3K9me3-based constitutive heterochromatin^11^. Both CENP-A and H3K9me3 serve as epigenetic templates for their inheritance through mitotic cell divisions via self-propagating epigenetic loops. CENP-A recruits the machinery for assembling new CENP-A nucleosomes^12^, and H3K9me3 recruits histone methyltransferases that catalyze formation of new H3K9me3^13^. Because these epigenetic mechanisms operate independently of DNA sequence, variation in satellite DNA composition was thought to have little or no functional impact^11^.

However, a key developmental stage in which pericentric satellites lack H3K9me3 occurs during early embryogenesis in various metazoans^14–20^. In a subset of these species, the lack of H3K9me3 specifically arises from the removal of histones from the paternal genome and their replacement by protamines during spermiogenesis^21–23^. CENP-A is an exception to this rule, as it is retained through spermiogenesis and thus serves as the epigenetic template for centromere inheritance through the male germline^24^. Following fertilization, maternal factors remove protamines and repackage the remainder of the paternal genome with histones initially lacking H3K9me3^25,26^, creating a unique developmental window in which paternal pericentric satellites are packaged by alternative forms of chromatin. Indeed, Polycomb Repressive Complexes 1 and 2 (PRC1 and PRC2) form what is typically considered facultative heterochromatin in place of H3K9me3 on paternal satellites in the mouse zygote^14,27,28^. Furthermore, PRC1 and PRC2 localize to chromatin by recognition of satellite DNA^27,28^, suggesting that natural variation in satellite sequence may alter the binding of these complexes and consequently influence heterochromatin formation and function.

In the widely used mammalian model species *Mus musculus* (house mouse), it takes approximately four cell cycles after fertilization for paternal satellites to gradually acquire H3K9me3 heterochromatin. Instead, PRC1 and PRC2 initially package paternal satellites in H2AK119ub1 and H3K27me3, respectively ^14,27,28^. Both modifications contribute to the transcriptional silencing of the 240 bp A/T-rich “major satellite” repeats at *M. musculus* pericentromeres^14,29^, with PRC1 also crucial for maintaining the integrity of these arrays^30^. Moreover, major satellite sequences can direct the recruitment of both PRC1 and PRC2. The Cbx2 subunit of PRC1 contains an AT-hook, a five amino acid motif with nanomolar binding affinity for the relatively narrow minor grooves of double stranded DNA (dsDNA) generated by stretches of tandem A/T nucleotides (A/T runs)^31–35^ (Fig 1A). It has therefore been proposed that PRC1 localization to paternal *M. musculus* pericentromeres is mediated by its AT-hook directly binding A/T runs present in major satellite repeats^28^. PRC2 localization to paternal pericentromeres likely requires BEND3, a methylation sensitive DNA-binding protein that can bind unmethylated major satellite DNA *in vitro*^27^.

**Figure 1.**
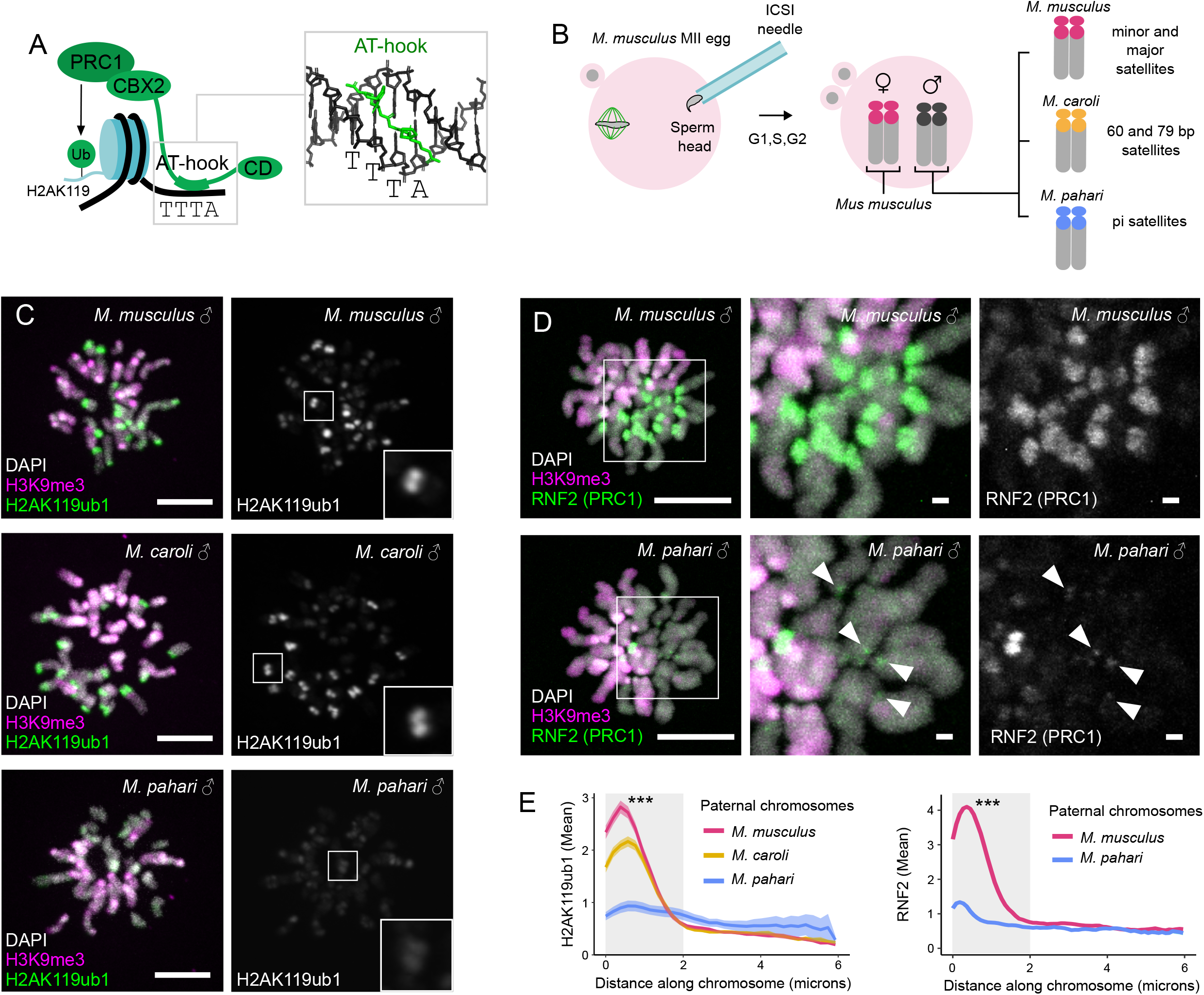
Natural variation in pericentric H2AK119ub1 heterochromatin and PRC1 recruitment. **A)** Schematic showing PRC1 recruitment to DNA via the AT-hook of its Cbx2 subunit, which binds into the minor groove of A/T runs present in satellite DNA. Chromodomain labelled as “CD”. **B)** Mouse zygotes are generated by injecting a *M. musculus* egg arrested in meiosis II with the sperm of mouse species with divergent pericentric satellite DNAs. **C-E)** Zygotes generated with sperm from the indicated species were arrested in mitosis with a kinesin-5 inhibitor (STLC, C) or by APC/C inhibition (proTAME, D), then fixed and stained for H2AK119ub1 or RNF2 (green), H3K9me3 (magenta) to mark maternal *M. musculus* chromosomes, and DAPI (gray). Arrowheads (D) highlight the slight enrichment of RNF2 near *M. pahari* centromeres. Graphs (E) show average H2AK119ub1 (n=103-164) and RNF2 (n=152-160) intensities along *M. musculus* (red), *M. caroli* (yellow) and *M. pahari* (blue) paternal chromosomes, starting from pericentric ends. S.E.M. is indicated by light band surrounding the mean line. Both *M. musculus* and *M. caroli* chromosomes have significantly higher H2AK119ub1 intensity than *M. pahari* chromosomes within the first two microns (gray box), (*** *P* <0.001). Statistical significance was calculated by a Kruskal-Wallis test, followed by Dunn’s post hoc test with Bonferroni correction. Images are max intensity z-projections, scale bars 10 μm or 1 μm (insets).

Here we ask whether natural variation in satellite sequences among closely related mouse species affects paternal pericentric heterochromatin in the zygote. By fertilizing *Mus musculus* eggs with the sperm of mouse species with divergent pericentric satellite DNA repeats, we find that *Mus musculus* pericentromeres form PRC1 heterochromatin whereas *Mus pahari* pericentromeres do not. AlphaFold3 modeling and DAPI binding indicate that the AT-hook of Cbx2 preferentially binds to specific A/T runs (i.e. AAAT) that form some of the narrowest minor grooves^36^, which are enriched in satellite arrays found at *M. musculus* but not *M. pahari* pericentromeres. We also find that PRC1 heterochromatin modifies pericentromere function by inhibiting the recruitment of the Chromosome Passenger Complex (CPC) to paternal *M. musculus* pericentromeres. The CPC regulates chromosome attachment to the mitotic spindle by destabilizing kinetochore-microtubule interactions. In zygotes, the distance between sister kinetochores in metaphase is larger for *M. musculus* vs *M. pahari* paternal chromosomes, indicating increased force pulling sister-kinetochores apart due to reduced CPC recruitment and stabilized kinetochore-microtubule interactions. Our results show that diverse pericentric DNA sequences lead to functionally diverse pericentromeres through differential binding of heterochromatin forming complexes.

## Results

### Variation in PRC1 localization and heterochromatin formation on divergent satellites

Given that major satellite DNA directs PRC1 and PRC2 heterochromatin formation, we hypothesized that divergent pericentric satellite sequences might alter Cbx2 or BEND3 binding and by extension H2AK119ub1 or H3K27me3 formation, respectively. To compare paternal chromosomes with different pericentric sequences, we developed an experimental system based on fertilizing *M. musculus* eggs with sperm from mouse species harboring genetically divergent pericentric satellites (Fig 1B). We selected three species (*M. musculus, M. caroli* and *M. pahari*) that differ in their satellite monomer sequence and not only in satellite copy number as seen in *M. musculus* subspecies and its sister-species *M. spretus*^2,37^. We injected sperm heads into MII eggs (i.e. intracytoplasmic sperm injection, ICSI^38^) to bypass reproductive barriers between species, arrested the resulting zygotes at the first mitosis by kinesin-5 inhibition, and measured H2AK119ub1 and H3K27me3 by immunofluorescence. Because only maternal chromosomes have H3K9me3 at this stage, we used this mark to distinguish maternal and paternal chromosomes^14^. Major transcriptional activation of the mouse embryo (zygotic genome activation) does not begin until the late two-cell stage^39^, so our analyses reflect interactions between satellite sequences and maternally inherited *M. musculus* PRC1/2 complexes rather than a mixture of subunits from both species. Analyzing heterochromatin in the zygote also allows us to test satellite DNA variation from evolutionarily distant species, which would otherwise be inaccessible due to hybrid incompatibilities that typically arise after zygotic genome activation^40–42^.

We found that all three species’ pericentromeres robustly acquire H3K27me3 (Fig S1). In contrast, H2AK119ub1 is enriched on paternal *M. musculus* and *M. caroli* pericentromeres relative to chromosome arms, but not on *M. pahari* pericentromeres (Fig 1C and 1E). To test whether these differences result from differences in PRC1 localization, we measured PRC1’s core-catalytic subunit, RNF2, by immunofluorescence. Consistent with the H2AK119ub1 enrichment, paternal *M. musculus* pericentromeres are strongly enriched for RNF2 relative to chromosome arms whereas *M. pahari* chromosomes only exhibit a slight enrichment near the very ends of their chromosomes, presumably near centromeres (Fig 1D and 1E). In certain contexts, PRC1 is recruited to chromatin by the chromodomain of its Cbx subunit, which directly binds H3K27me3^43–45^, and Cbx2 also contains a chromodomain (Fig 1A). However, the presence of H3K27me3 at *M. pahari* paternal pericentromeres (Fig S1), which lack H2AK119ub1 and RNF2 (Fig 1C-E), indicates that H3K27me3 is not sufficient for PRC1 recruitment during the first zygotic mitosis, consistent with previous findings that PRC1 can localize to paternal pericentromeres independent of PRC2 and H3K27me3^14^. Overall, these results demonstrate striking differences in PRC1 recruitment and heterochromatin formation at divergent pericentric satellites, independent of PRC2.

### Satellite sequence determinants of PRC1 binding

To explain the differences in pericentric RNF2 and H2AK119ub1, we hypothesized that *M. musculus* and *M. caroli* pericentromeric satellites, but not *M. pahari* satellites, are enriched for narrow DNA minor grooves that bind the AT-hook of Cbx2 to recruit the PRC1 complex. Following a previously described approach^2^, we used DAPI staining to compare the amount of narrow minor grooves generated by A/T-runs at each species’ pericentromere, as DAPI specifically binds to these regions on the dsDNA molecule^46–48^. Consistent with previous observations^2,9,49^, we find increased DAPI staining at *M. musculus* and *M. caroli* pericentromeres relative to chromosome arms, but not at *M. pahari* pericentromeres (Figure 2A, 2B and Fig S2A). To exclude the possibility that DAPI enrichment at pericentromeres represents a relatively more compact chromatin state compared to the rest of chromosome, we stained with Sytox Green, an intercalating DNA-dye with no considerable DNA sequence preference^50^. None of the three species show enrichment of Sytox Green at their pericentromeres (Fig 2C, 2D and Fig S2B). Together, these results demonstrate that *M. musculus* and *M. caroli* pericentromeres are enriched for narrow A/T-rich minor grooves that can robustly bind the AT-hook of Cbx2, whereas *M. pahari* pericentromeres are not.

**Figure 2.**
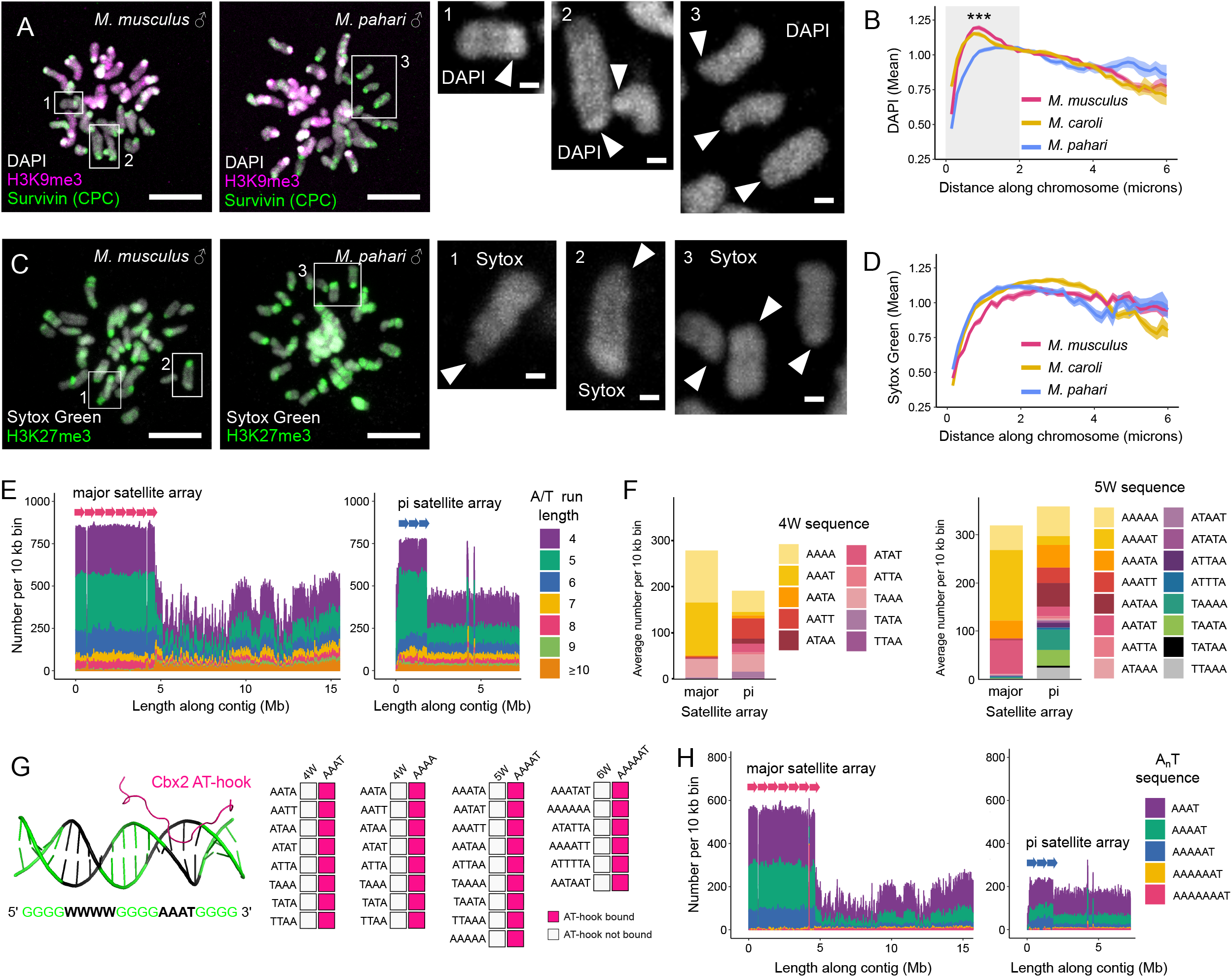
*M. musculus* major satellite arrays are enriched for A_n_T sequences that preferentially bind the Cbx2 AT-hook. **A-D)** Zygotes generated with *M. musculus* or *M. pahari* sperm were arrested in mitosis with a kinesin-5 inhibitor (STLC), then fixed and stained as indicated. Survivin (CPC subunit) marks pericentromeres, H3K9me3 marks maternal chromosomes, H3K27me3 mark paternal pericentromeres, and DAPI or Sytox Green label DNA. Arrowheads in insets point to paternal pericentromeres as indicated by presence of Survivin and absence of H3K9me3 (A) or enrichment of H3K27me3 (C). Images are max intensity z-projections, scale bars 10 μm or 1 μm (insets). Graphs show average DAPI (B, n=72-165) or Sytox Green (D, n=44-66) intensity along *M. musculus* (red), *M. caroli* (yellow) and *M. pahari* (blue) paternal chromosomes, starting from pericentric ends. Both *M. musculus* and *M. caroli* chromosomes have significantly higher DAPI intensity than *M. pahari* chromosomes within the first two microns (B, gray box), (*** *P* <0.001). Statistical significance was calculated by a Kruskal-Wallis test, followed by Dunn’s post hoc test with Bonferroni correction. **E)** Histograms show the number of various A/T run lengths per 10 kb bin along portions of two genomic contigs containing arrays of either *M. musculus* major satellite (left) or *M. pahari* pi satellite (right), indicated by tandem arrows. **F)** The average number per 10 kb bin of all possible 4W (left) and 5W (right) sequences within the major and pi satellites arrays shown in E. **G)** Schematic (left) shows *in silico* competition assays between different A/T run sequences for the binding of a single peptide encoding the AT-hook of Cbx2, using AlphaFold3. Results (right) show AT-hook binding preferences. **H)** Histograms show the number of various A_n_T sequences per 10 kb bin along the same genomic contigs as in E.

We considered two possibilities for the underlying differences in pericentric DNA sequence between species, leading to differences in DAPI enrichment and PRC1 localization. We focused on comparisons between *M. musculus* and *M. pahari* because of the long-read sequencing data available for these species. First, *M. musculus* major satellites may have a higher frequency of A/T runs compared to *M. pahari* pericentric satellites (pi satellites)^9^. However, major and pi satellite consensus sequences have similar frequencies of A/T runs of lengths ranging from four (4W, the minimal number of nucleotides the AT-hook can span) to ten or more (≥10W) (Fig S2C). To capture nucleotide diversity that may be lost in consensus sequences, we also calculated the frequencies of A/T runs across genomic contigs containing either major or pi satellite arrays. Consistent with consensus sequences, we find that both species’ satellite arrays have similar frequencies of A/T runs (∼825 vs ∼750 per 10 kb window in major satellite or pi satellite, respectively) (Fig 2E). The frequencies are also similar for each length of A/T runs (Fig 2E and Fig S2D), with the exceptions of 6W and ≥10W, which are relatively more abundant in major and pi satellite arrays, respectively. However, it is unclear how differences in these lengths specifically would lead to higher PRC1 and DAPI at *M. musculus* major satellite arrays. Therefore, it is unlikely that differences in A/T-run frequency explain the difference in PRC1 heterochromatin formation.

The second possibility is that *M. musculus* major satellite arrays might be enriched for specific A/T sequences optimal for Cbx2 AT-hook binding, compared to *M. pahari* satellite arrays. We analyzed the sequence composition of 4W, 5W and 6W A/T runs, which make up ∼75% of the A/T stretches in both species’ satellite arrays and therefore most potential AT-hook binding sites. The vast majority of 4W sequences in *M. musculus* major satellite arrays are AAAT (114 per 10 kb), AAAA (113 per 10 kb) or TAAA (39 per 10 kb) (Fig 2F and Fig S2E). *M. pahari* pi satellite arrays have a similar frequency of TAAA sites (37 per 10 kb) but much lower frequencies of AAAT and AAAA sites (7.2 and 46 per 10 kb, respectively). Based on these differences, we hypothesized that the Cbx2 AT-hook preferentially binds AAAT and/or AAAA.

To test the binding preferences of the Cbx2 AT-hook, we used AlphaFold3^51^ to perform *in silico* pairwise competitive binding assays for AAAT and AAAA vs all other possible A/T tetranucleotides (Fig 2G). We first confirmed that AlphaFold3 accurately models AT-hook binding to dsDNA using the well characterized HMGA1 AT-hook bound to a fragment of the IFN-ß promoter^34^ (PDB ID 2EZD) as a reference. The predicted structure closely matched the known structure (RMSD=0.48 Å), and in both structures, the AT-hook occupied the minor groove of the A/T run (AAATT) (Fig S2F). Furthermore, the confidence scores of the predicted structure (iPTM=0.48 and PTM=0.47) are similar to the models generated in our *in silico* competitive binding assay (Table S1). The AT-hooks of Cbx2 and HMGA1 share the same core sequence (PRGRP) which binds the minor groove, supporting our use of AlphaFold3 to model the AT-hook binding preference of Cbx2.

For each competition assay, we model a single dsDNA molecule encoding two distinct A/T runs flanked by four G:C base pairs (5’ GGGGWWWWGGGGWWWWGGGG 3’). These two A/T runs “compete” to bind a single peptide consisting of the core Cbx2 AT-hook along with flanking amino acids (RKRGKR**PRGRP**RKHTVTSS). All AlphaFold models predict that the Cbx2 AT-hook prefers to bind AAAT and AAAA over all other A/T tetranucleotides (Fig 2G). When AAAT and AAAA compete against each other, their relative order on the dsDNA dictates which sequence binds the AT-hook (Fig S2G, see Materials and Methods), indicating that these two sequences have similar affinity for the AT-hook. We obtain similar results when analyzing 5W and 6W A/T runs. AAAAT and AAAAAT are the most abundant 5W and 6W sequences in the *M. musculus* major satellite array (147 per 10 kb and 70 per 10 kb, respectively) (Fig 2F and Fig S2E). These sequences outcompete all other prevalent 5W and 6W sequences in both major and pi satellite arrays for Cbx2 AT-hook binding (Fig 2G and Fig S2G). Furthermore, AAAAT and AAAAAT are 8 and 2.5-fold less frequent, respectively, in *M. pahari* pi satellite compared to *M. musculus* major satellite arrays (Fig 2F and Fig S2E).

These results indicate that the Cbx2 AT-hook prefers to bind stretches of adenine nucleotides that end with a single thymine (A_n_T), consistent with previous reports showing that the core motif of the AT-hook (PRGRP) preferentially binds the minor groove of the sequence 5’ AA(A/T)T 3’. This preference reflects optimal Van der Waals packing when adenine bases are on opposite sides of the A/T run^35^. Our AlphaFold3 models also show that the AT-hook, which can span 4-5 bases, spans AAAT or AAAAT bases within all but one modeled AAAAAT sequence, rather than the overlapping AAAA or AAAAA bases (Fig S2H and Table S1).

Additionally, a meta-analysis of PDB structures^36^ indicates that AAAT generates one of the narrowest minor grooves compared to other tetranucleotide sequences, which may contribute to preferential binding of both the AT-hook and DAPI. To generalize this analysis to include A_n_T sequences (like AAAT) present in longer A/T runs, we quantified the frequency of A_n_T sequences of various lengths (i.e. AAAT, AAAAT, AAAAAT, AAAAAAT and AAAAAAAT) in major and pi satellite arrays independent of total A/T run length. We find that major satellite arrays have approximately three times more of these A_n_T sequences than *M. pahari* pi satellite arrays (Fig 2H and Fig S2I). Furthermore, pi satellite arrays exhibit A_n_T frequencies similar to euchromatin. Overall, these analyses indicate that PRC1 and its H2AK119ub1 modification are enriched at *M. musculus* major satellite arrays because they are enriched for A_n_T sequences that preferentially bind the Cbx2 AT-hook. In contrast, *M. pahari* pi satellite arrays exhibit a lower frequency of A_n_T sequences like euchromatin and fail to enrich both PRC1 and H2AK119ub1.

### Consequences of PRC1 heterochromatin for pericentromere function

Next, we asked if the difference in PRC1-heterochromatin between paternal *M. musculus* and *M. pahari* chromosomes has consequences for pericentromere function. Notably, H2AK119ub1 is positioned close to the site of another post-translational modification: phosphorylation of H2AT121 by Bub1 kinase. H2AT121phos recruits the Chromosome Passenger Complex (CPC) to pericentromeres via Sgo1 (Fig 3A) to ensure accurate chromosome segregation by regulating kinetochore-microtubule interactions during mitosis^52– 54^. ChIP-seq and *in vitro* assays on reconstituted nucleosomes have shown that H2AK119ub1 and H2AT121phos are mutually exclusive^55^, suggesting that one modification physically blocks the deposition of the other on the same H2A C-terminal tail. We hypothesized that H2AK119ub1 occludes Bub1 kinase from phosphorylating H2AT121 and thereby reduces CPC binding. This hypothesis predicts low H2AT121phos and low CPC on paternal *M. musculus* pericentromeres (Fig 3A), which have high H2AK119ub1 compared to maternal *M. musculus* and paternal *M. pahari* pericentromeres (Fig 1C and 1E).

**Figure 3.**
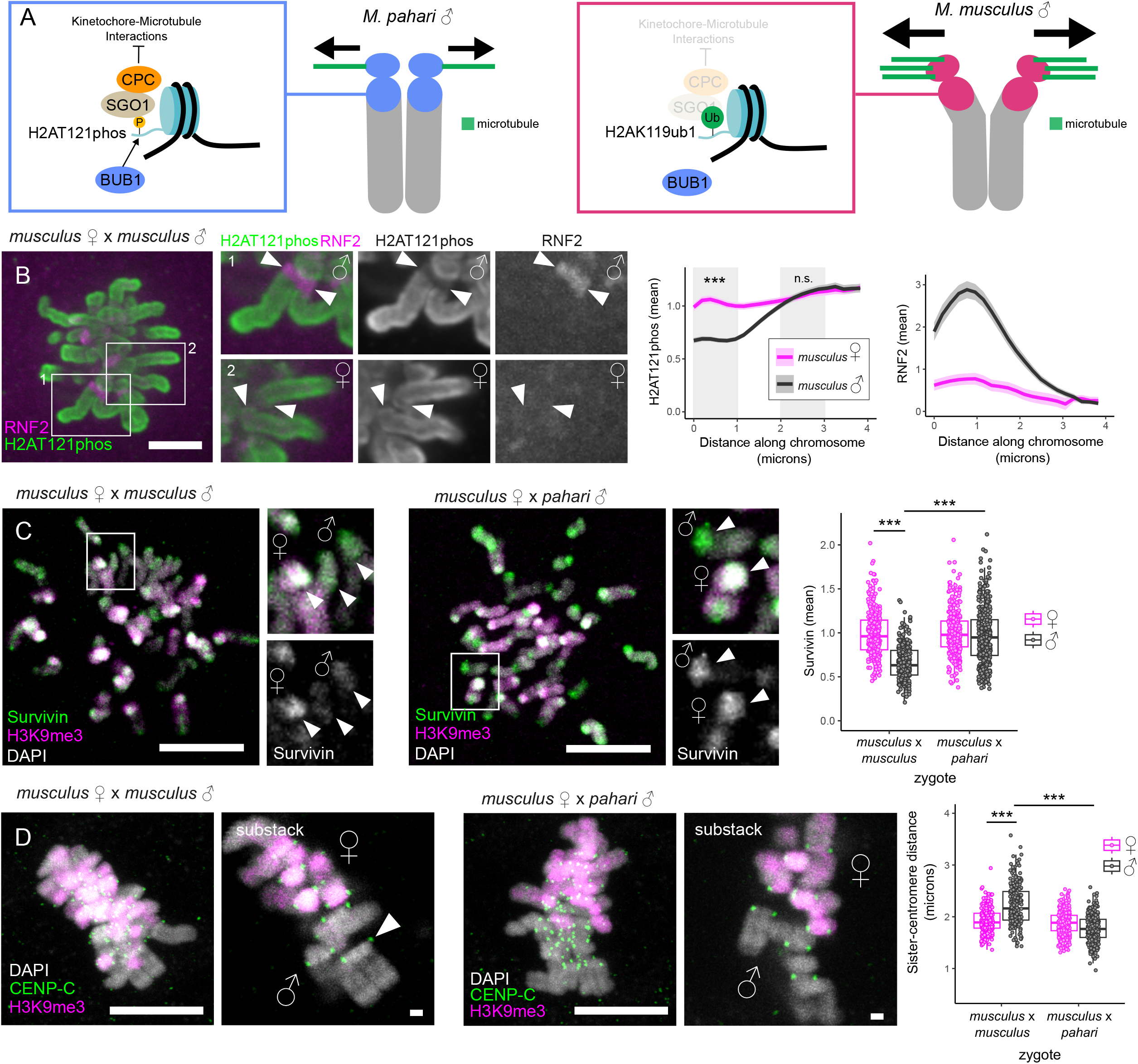
PRC1 pericentric heterochromatin inhibits CPC recruitment. **A)** Schematic shows how H2AK119ub1 can inhibit H2AT121phos and CPC recruitment, resulting in increased distance between metaphase sister kinetochores at paternal *M. musculus* pericentromeres (right) vs *M. pahari* pericentromeres (left). **B)** Pure *M. musculus* zygotes were arrested at metaphase by APC/C inhibition (proTAME), then fixed and stained for RNF2 (PRC1 subunit, magenta) and H2AT121phos (green). Top and bottom insets show paternal (RNF2 positive) and maternal (RNF2 negative) chromosomes, respectively; arrowheads point to pericentromeric ends of chromosomes. Graphs show average H2AT121phos and RNF2 intensity along maternal (n=56) and paternal (n=54) chromosomes, starting from pericentric ends. Statistical differences between maternal and paternal chromosomes from 0-1 and 2-3 microns along their length (gray boxes) were calculated. **C)** Zygotes generated with *M. musculus* or *M. pahari* sperm were arrested in mitosis with a kinesin-5 inhibitor (STLC), then fixed and stained for Survivin (green) and H3K9me3 (magenta). Arrowheads in insets point to pericentric Survivin staining. Graph shows the mean Survivin intensity of maternal and paternal pericentromeres in both types of zygotes. Each point represents a single pericentromere (n=253-374 for each group) and boxes represent interquartile ranges. **D)** Zygotes generated with *M. musculus* or *M. pahari* sperm were arrested at metaphase by APC/C inhibition (proTAME), then fixed and stained for CENP-C (green) and H3K9me3 (magenta). Arrowhead points to a pair of paternal *M. musculus* sister kinetochores. Graph shows distances between sister kinetochores for maternal and paternal chromosomes in both types of zygotes. Each point represents a single pair (n=230-284 for each group). All *P*-values were calculated by a Kruskal-Wallis test followed by Dunn’s test using Bonferroni correction for multiple testing (*** *P* < 0.001). Images are max intensity z-projections; scale bars 10 μm or 1 μm (insets).

To test this prediction, we first visualized H2AT121phos, the CPC subunit Survivin, and the PRC1 subunit RNF2. In *M. musculus* ♀ x *M. musculus* ♂ zygotes arrested at metaphase by chemical inhibition of the anaphase promoting complex/cyclosome (APC/C), we found that H2AT121phos is present along the entire chromosome (Fig 3B) as reported in the previous cell cycle in mouse oocytes^56,57^. Moreover, we found an inverse relationship between RNF2 and H2AT121phos, with the presence of RNF2 at paternal pericentromeres coinciding with a reduction in H2AT121phos (Fig 3B). In contrast, maternal pericentromeres with undetectable RNF2 show no reduction in H2AT121phos. Consistently, when we compared regions of maternal and paternal chromosomes that both lack RNF2, H2AT121phos intensities were similar, indicating that reduced H2AT121phos depends on RNF2. On average, H2AT121phos is reduced to ∼60% at paternal *M. musculus* pericentromeres compared to maternal pericentromeres (Fig 3B). Consistent with paternal *M. musculus* H2AK119ub1 heterochromatin inhibiting H2AT121phos, Survivin intensity on paternal *M. musculus* pericentromeres is reduced to ∼65% compared to paternal *M. pahari* and maternal *M. musculus* pericentromeres, both of which form relatively less H2AK119ub1 (Fig 3C). Together, these results show that PRC1 heterochromatin inhibits the acquisition of H2AT121phos and thereby reduces CPC recruitment to pericentromeres of paternal *M. musculus* chromosomes.

The Aurora B kinase subunit of the CPC phosphorylates kinetochore proteins to destabilize kinetochore-microtubule interactions^58^. In tissue culture cells, expression of a non-phosphorylatable mutant of a key Aurora B substrate, Hec1, results in increased stabilization of kinetochore-microtubule interactions and increased forces pulling the sister kinetochores apart at metaphase^59^. Increased inter-kinetochore distance therefore provides a functional readout for reduced Aurora B activity (Fig 3A). To test for functional consequences of CPC differences between pericentromeres in the zygote, we measured the distance between sister kinetochores at metaphase. We found increased inter-kinetochore distances for paternal *M. musculus* chromosomes compared to either maternal *M. musculus* or paternal *M. pahari* chromosomes (Fig 3D). This result is consistent with our hypothesis that PRC1 heterochromatin inhibits H2AT121 phosphorylation, downregulates the CPC, and stabilizes kinetochore-microtubule interactions on paternal *M. musculus* chromosomes. Increased inter-kinetochore distances might also reflect reduced pericentric cohesion because in addition to recruiting the CPC, Sgo1 also recruits phosphatases that counteract cohesin removal that occurs along chromosome arms during mitotic prophase^60,61^ (Fig 3A). However, sister chromatids appear in close proximity along most of their lengths during zygotic mitosis^62^ (Fig 3D), suggesting that cohesins are not removed during prophase at this stage of development. Therefore, we interpret the increase in inter-kinetochore distance as a consequence of reduced CPC activity.

## Discussion

Long-read sequencing technology continues to reveal the extensive diversity of satellite DNA sequence composition^8,9,63^, but experimental systems that test the functional impact of this diversity remain limited. In the mouse model species *M. musculus*, centromeric and pericentromeric satellite variation is limited to copy-number differences of the same satellite sequences^2,37,56^. Our hybrid embryo system overcomes this limitation by incorporating sequence variation between satellites that is only found in more divergent non-model mouse species. Using this system, we demonstrated that *M. musculus* major satellite sequence is more effective than *M. pahari* pi satellite sequence at recruiting PRC1. AlphaFold3 modeling indicates that this difference results from the AT-hook of Cbx2^*M*. *musculus*^ preferentially binding the minor groove of A_n_T sequences, which are enriched in *M. musculus* major satellite arrays when compared to *M. pahari* pi satellite arrays and the rest of the genome. The sequence preference for the Cbx2 AT-hook is consistent with the known optimal binding sequence of other AT-hook proteins^35^. We propose that the relatively high density of A_n_T sequences at *M. musculus* major satellite arrays is above a threshold required for efficient PRC1 recruitment and robust H2AK119ub1 formation.

Previous findings have shown that the Cbx2 chromodomain, in addition to the AT-hook, contributes to PRC1 localization to chromatin during zygotic interphase, likely by binding H3K27me3 that first forms during S-phase^28^. However, our data demonstrate that during mitosis, paternal *M. pahari* pericentromeres exhibit similar levels of H3K27me3 as paternal *M. musculus* pericentromeres (Fig S1A) yet significantly reduced PRC1 and H2AK119ub1 (Figure 1C-E). These findings indicate that the Cbx2 chromodomain does not effectively localize PRC1 during mitosis. We speculate that Aurora B-mediated phosphorylation of H3S28 during mitosis^64^ prevents the Cbx2 chromodomain from binding H3K27me3, similar to how Aurora B-mediated phosphorylation of H3S10 inhibits HP1 chromodomain binding to H3K9me3^65^. As a result, satellite DNA binding via the Cbx2 AT-hook predominately dictates pericentric PRC1 localization and H2AK119ub1 heterochromatin during mitosis. Nonetheless, during interphase, the Cbx2 chromodomain likely binds H3K27me3 at *M. pahari* satellites, recruiting PRC1 and establishing H2AK119ub1. Upon mitotic entry, however, PRC1 is displaced from *M. pahari* chromatin and cannot counteract the removal of pericentric H2AK119ub1 by cytosolic deubiquitinating enzymes once the nuclear envelope breaks down^66^. Thus, our analysis of mitotic chromosomes indicates that Cbx2’s chromodomain is not required at this phase of the cell cycle, highlighting cell-cycle-specific differences in how PRC1 is recruited to chromatin.

Despite the importance of PRC1 pericentric heterochromatin in *M. musculus* zygotes^30^, our findings suggest that it is not a conserved feature of mitotic paternal pericentromeres, as it is lacking from *M. pahari* pericentromeres. *M. musculus* major satellites and *M. caroli* satellites may have specific properties that require their packaging in PRC1 heterochromatin, whereas *M. pahari* pi satellites can rely on PRC2-mediated H3K27me3. Alternatively, the *M. pahari* egg cytoplasm may have specific adaptations that allow PRC1 to be recruited to mitotic pericentromeres by a different mechanism. Because the core Cbx2 AT-hook sequence is conserved across Muridae (data not shown), it is unlikely that the AT-hook of Cbx2^*M*. *pahari*^ has adapted to bind A/T sequences enriched at *M. pahari* pi satellites. Similarly, the Cbx2 chromodomain is identical across Muridae (data not shown), indicating that the chromodomain of Cbx2^*M*. *pahari*^ has not adapted to have a higher binding affinity towards H3K27me3. However, vertebrates encode multiple PRC1 complexes that share the core ubiquitin ligase subunits (RING1A/1B) but differ in associated proteins that diversify PRC1 function, including how the complex is recruited to chromatin^67^. Thus, another chromatin targeting subunit expressed in *M. pahari* zygotes might recognize paternal *M. pahari* pericentromeres during mitosis.

Although maternal and paternal major satellites are genetically identical, only paternal satellites robustly form PRC1 heterochromatin because H3K9me3-based heterochromatin on maternal satellites inhibits PRC1 recruitment^14,28^. Given this, we propose that in most cell cycles H3K9me3 can serve as a buffer against DNA sequence-based pathways of heterochromatin formation, suppressing functional diversity that might otherwise arise from differences in satellite sequences. In contrast, our findings highlight the first zygotic cell cycle as a unique environment where PRC1 heterochromatin formation and function are sensitive to variation in satellite DNA sequence due to the loss of H3K9me3 from the paternal genome during spermiogenesis. This functional variation occurs at a critical stage of development where chromosome segregation errors would lead to aneuploidy of all subsequent cells.

Despite the importance of this first cell division, PRC1 heterochromatin inhibits molecular pathways that ensure accurate chromosome segregation, particularly regulation of kinetochore-microtubule attachments by the CPC. Our findings suggest that expansion of pericentric satellite arrays enriched for A_n_T sequences would lead to increased PRC1 heterochromatin and a corresponding decline in CPC recruitment, making paternal chromosomes more prone to segregation errors. Although it is difficult to test for potentially subtle differences in error rates between paternal and maternal chromosomes in mouse zygotes, our hypothesis predicts selection against large major satellite arrays in natural populations. Consistent with this idea, wild-caught *M. musculus* have roughly ten times fewer major satellite repeats than inbred laboratory strains, where natural selection is relatively weak^37^. Despite potentially deleterious effects, however, pericentric PRC1 heterochromatin is required for the transcriptional silencing and stability of paternal major satellite arrays^28,30^, suggesting an evolutionary trade-off between proper packaging of major satellites and chromosome segregation fidelity.

## Materials and Methods

### Mouse strains

The mouse strain representing *Mus musculus* in our experiments is FVB/NJ, purchased from Jackson Laboratory (strain#001800). FVB/NJ was chosen because its MII eggs can survive intracytoplasmic sperm injection and because it is an inbred strain. *M. caroli* (CAROLI/EiJ, strain#000926) and *M. pahari* (PAHARI/EiJ, strain #002655) were also purchased from Jackson Laboratory. We maintain our own colony of *M. pahari* because Jackson Laboratory no longer carries this species. All animal experiments and protocols were approved by the Institutional Animal Use and Care Committee of the University of Pennsylvania and are consistent with National Institutes of Health guidelines (protocol #804882).

### Intracytoplasmic sperm injection (ICSI)

To prepare sperm for ICSI, the epididymis and vas deferens were removed from males and dissected in PBS to release mature sperm. Sperm were allowed to swim-out for 10 mins on a 37°C slide warmer. After swim out, sperm were pelleted by centrifugation at 700xg at 4°C for 5 min. The PBS supernatant was removed, and sperm were then washed twice with ice-cold Nuclear Isolation Media (NIM, 123 mM KCl, 2.6mM NaCl, 7.8 mM Na_2_PO_4,_ 1.4 mM KH_2_PO_4_, 3 mM EDTA, pH adjusted to 7.2 using 1M KOH) with 1% poly vinyl alcohol (PVA). After the second wash, sperm were resuspended in 100 μL of NIM 1% PVA and then sonicated using a Branson Sonic Bath Model 1210 for 15-20 second intervals until at most 30% of sperm had their heads detached from their tails. Sonicated sperm were washed twice with NIM 1% PVA. After the second wash, the sperm pellet was resuspended in a 1:1 solution of glycerol and NIM 1% PVA and placed at -20°C until the day of injection (the following day or 1 week later).

Females were super-ovulated by injection with 5 U PMSG (Peptides International) followed by injection with 5 U hCG (Sigma) 48 hours later. MII eggs were collected from females 14-15 hours post-hCG injection and placed in M2 media (Sigma) with hyaluronidase (0.15 mg/mL) to remove cumulus cells. MII eggs were then washed through four drops of M2 media supplemented with 4 mg/mL BSA (M2+BSA) and left on a 37°C plate warmer until injection. 10-20 μLs of sonicated sperm was diluted in 200 μL of NIM 1% PVA, gently vortexed, and then washed twice with NIM 1% PVA. After the last wash, the sperm pellet was resuspended in 100 μL of NIM 1% PVA and left on ice until injection.

Batches of 10 MII eggs were injected with sperm heads at a time in room temperature drops of M2+BSA. After injection, each batch was moved to a drop of M2+BSA on a 37°C plate warmer and allowed to recover for 1 hour. After recovery, each batch was washed through four drops of pre-equilibrated AKSOM (Millipore Sigma) and placed in at 37°C humidified incubator with 5% CO_2_ until injections were complete. Depending on the experiment, at least three hours after injection, eggs were then moved into a drop of AKSOM with either 10 μM STLC (kinesin-5 inhibitor, Sigma) (to arrest cells in mitosis with monopolar spindles) or 5 μM proTAME (APC/C inhibitor, R&D Systems) (to arrest zygotes in metaphase), then allowed to develop overnight in a 37°C incubator with 5% CO_2_.

### Fixing and staining

The morning after ICSI, arrested zygotes were fixed in 37°C 2% paraformaldehyde in PBS for 20 minutes, washed through three pools of blocking solution (PBS containing 0.5% BSA, 0.01% Tween-20) and then left at 4°C overnight. The next morning cells were permeabilized in PBS containing 0.5% Triton X-100 (Sigma) for 15 minutes at room temperature, quickly washed through two pools of blocking solution and then allowed to block for 20 minutes at room temperature. For H3K9me3 and H3K27me3 staining, the cells were treated with lambda-phosphatase (1600 U, NEB) for 1 hour at 37°C in a humidified chamber. Embryos were then quickly washed through two pools of blocking solution and then incubated with primary antibodies for 1 hour in a dark humidified chamber at room temperature (or overnight at 4°C for H2AT121phos staining). Afterwards, embryos underwent three 15-minute washes in blocking solution and then were incubated with secondary antibodies for 1 hour in a dark humidified chamber at room temperature. Cells then underwent three 15-minute washes in blocking solution and were mounted in Vectashield with DAPI (Vector) to stain chromosomes. For cells that were stained with Sytox Green (Invitrogen), the first wash after incubation with secondary antibodies included 1 uM Sytox Green and after the next two washes were mounted in Vectashield without DAPI. Primary antibodies used are rabbit anti-H2AK119ub1 (1:800, Cell Signaling, D27C4), rabbit anti-H3K27me3 (1:700, Cell Signaling, C36B11), mouse anti-H3K9me3 (1:200, Active Motif, 39285), rabbit anti-H3K9me3 (used when co-staining against RNF2, 1:500, Active Motif, 39162), rabbit anti-Survivin (1:500, Cell Signaling, 71G4B7), rabbit anti-H2AT120phos (1:2500, Active Motif, 39392), mouse anti-RNF2/Ring1B (1:500, Active Motif, 39664) and rabbit anti-CENP-C^68^ (1:500). Secondary antibodies used are donkey anti-rabbit Alexa Fluor 488, donkey anti-mouse Alexa Fluor 594, donkey anti-rabbit Alexa Fluor 594, and donkey anti-mouse Alex Fluor 647. All secondary antibodies are purchased from Invitrogen and used at a dilution of 1:500.

### Microscopy

Some confocal images were collected as z-stacks with 0.5 μm intervals, using a microscope (DMI4000 B; Leica) equipped with a 63× 1.3 NA glycerol-immersion objective lens, an xy piezo Z stage (Applied Scientific Instrumentation), a spinning disk confocal scanner (Yokogawa Corporation of America), an electron multiplier charge-coupled device camera (ImageEM C9100–13; Hamamatsu Photonics), and either an LMM5 (Spectral Applied Research) or Versalase (Vortran Laser Technology) laser merge module, controlled by MetaMorph software (Molecular Devices, v7.10.3.294). Confocal images were also collected using a Leica TCS SP8 Four Channel Spectral Confocal System with a 63x objective lens. A z-stack of 0.3 μm intervals was used to collect images of embryos used to calculate sister-kinetochore distances. All samples in an experiment were imaged using the same laser settings.

### Image quantification and statistical analysis

All image analysis was carried out using ImageJ/Fiji^69^. To measure signal intensities along single chromosomes, we selected chromosomes that were mostly in the XY plane and generated max intensities projections of them. Beginning from the pericentric end, we then used the segmented line tool to manually draw a linear ROI (roughly the width of the chromosome) along the chromosome’s length and measured the average signal intensity per unit distance along the ROI. Average background was then subtracted from these values. To more easily compare the distributions of DAPI and Sytox Green intensity, their values were normalized to the average intensity within each cell/embryo using custom R scripts.

To measure Survivin intensity at pericentromere ends, we also selected chromosomes that were mostly in the XY plane and generated maximum intensity projections of them. We then use the rectangular selection tool to generate square ROIs that encompassed pericentric ends of chromosomes and measured average intensity within the ROI. Average background was then subtracted from these values. The resulting values were then normalized to maternal chromosomes in each cell/embryo using custom R scripts.

To measure the distance between sister-centromeres, we uniquely labelled each centromere and extracted its 3D coordinates using 3D maxima finder and 3D Roi Manager (based on CENP-C signal). We then manually assigned sister-centromeres with the help of BigDataViewer to “rotate” the image stack in cases where the spindle pole axis was not in the XY plane and then used custom R scripts to calculate the distance between them.

*P*-values were determined using R. We performed non-parametric Kruskal-Wallis tests followed by Dunn’s test and used the Bonferroni method to correct *P-*values for multiple testing. Plots were made in the ggplot2 package in R.

### Satellite sequence analysis

*M. musculus* assemblies containing major satellite arrays were found by BLASTing the major satellite consensus sequence^37^ on NCBI BLAST server. These assemblies are from a long-read *M. musculus* genomes generated by the Mouse Genome Project and Darwin Tree of Life Project (https://www.nhm.ac.uk/our-science/research/projects/darwin-tree-of-life.html#:~:text=The%20Darwin%20Tree%20of%20Life%20project%20aims%20to%20generate%20DNA,fungi%20within%20the%20British%20Isles.). The pi satellite array containing contigs are from a published *M. pahari* long-read assembly^9^. These contigs were chosen because they represent relatively complete assemblies of functional *M. pahari* centromeres and pericentromeres based on CENP-A and H3K9me3 ChIP-seq. We chose to study major and pi satellite arrays because they are the most abundant pericentric satellites and because PRC1 is known to associate with major satellite^14^.

To quantify the frequency of A/T runs of various lengths along a contig, we used the matchPattern() function in the Biostrings package in R (https://bioconductor.org/packages/Biostrings). Specifically, we searched for A/T runs that were at least four nucleotides long (i.e. matched “WWWW”), with overlapping matches eventually being merged into one longer A/T run by using the reduce() function. To analyze the composition of 4W and 5W A/T runs, we used the matchPattern() function to find all possible 4W and 5W sequences and then only kept matches that overlapped with 4W or 5W A/T run lengths, respectively. To analyze the composition of 6W A/T runs, we used the 6W A/T run coordinates to extract sequences within the satellite array using the Genomic Ranges package^70^ in R. Sequences that are reverse complements of each other are redundant and therefore were merged into one. The average number of times a particular sequence occurred per 10 kb bin within an array was calculated by dividing the total number of occurrences of a particular sequence within the array by the length of the array and then multiplying by 10,000. We defined the range of the satellite arrays for each contig based on coordinates from satellite consensus sequence BLAST results. Finally A_n_T sequences were also found along contigs using the matchPattern() function. To prevent over-counting of smaller AnT sequences (e.g. overlap of AAAT and AAAAT), only the longest of overlapping A_n_T sequences were counted.

### AlphaFold3 modeling

All sequences were modeled using the AlphaFold3 webserver (https://alphafoldserver.com/). To model a single dsDNA molecule encoding two distinct A/T runs flanked by four G:C base pairs, we entered a single forward and a single reverse strand (e.g. forward: 5’GGGGAAAAGGGGAAATGGGG 3’, reverse: 5’CCCCATTTCCCCTTTTCCCC 3’). We also modeled a single peptide encoding the AT-hook of Cbx2 with flanking amino acids (RKRGKR**PRGRP**RKHTVTSS). Alphafold3 outputs five model structures (0-4), with 0 scored as the best and the fourth scored as the worst. Each model was visually inspected using Pymol (The Pymol Molecular Graphics System, Version 3.0 Schrodinger, LLC). All five models for each competition assay were consistent with one another (Table S1). To rule out the possibility that the relative position along the dsDNA molecule dictates where the AT-hook binds, we flipped the relative positions of the two A/T sequences along the dsDNA. For example, we modeled both 5’GGGGAAAAGGGGAAATGGGG 3’ and 5’GGGGAAATGGGGAAAAGGGG3’ sequences (Figure S2G).

## Acknowledgements

We thank Dr. Colin Conine for training in intracytoplasmic sperm injection and Dr. Ben Black and Dr. Damian Dudka for thoughtful conversations on AT-hook proteins. We also thank Dr. Black for initially suggesting that pericentric H2AK119ub1 may be incompatible with H2AT121phos. We thank Dr. Beth L. Dumont for sending us PAHARI/EiJ mice from Jackson Laboratory. We thank Emma Hamlin for help with data analysis. We thank Drs. Mia Levine, Ben Black and Richard Schultz for helpful discussions and comments on the manuscript.

## Author Contributions

Conceptualization: P. Lamelza and M. A. Lampson, Methodology: P. Lamelza and M. Parrado, Investigation: P. Lamelza, and M. Parrado, Writing: P. Lamelza and M. A. Lampson, Funding Acquisition: P. Lamelza and M.A. Lampson, Resources: M.A. Lampson

