## Supplemental Figures for "Species-specific satellite DNA composition dictates PRC1-mediated pericentric heterochromatin"

**Figure S1. Paternal *M. musculus, M. caroli* and *M. pahari* chromosomes acquire pericentric H3K27me3. A)** Zygotes generated with sperm from the indicated species were arrested in mitosis using a kinesin-5 inhibitor, then fixed and stained for H3K27me3 (green), H3K9me3 (magenta) to mark maternal *M. musculus* chromosomes, and DAPI (gray). Graphs plot average H3K27me3 intensities along *M. musculus* (red, n=99), *M. caroli* (yellow, n=135) and *M. pahari* (blue, n=130) paternal chromosomes, starting from pericentric ends. S.E.M. is indicated by light band surrounding the mean line.

**Figure S2. Satellite sequence determinants of PRC1 binding. A-B)** Zygotes generated with *M. caroli* sperm were arrested in mitosis with a kinesin-5 inhibitor (STLC), then fixed and stained for H3K27me3 (green) to mark paternal pericentromeres and either DAPI (A, gray) or Sytox Green (B, gray). Arrowheads in insets point at paternal *M. caroli* pericentromeres. Images are max intensity z-projections; scale bars 10 μm or 1 μm (insets). **C)** Major and pi satellite consensus sequences (left) with A/T runs greater than or equal to four shown in bold. Table (right) summarizes the number of specific A/T run lengths (4W to $\geq$10W) in satellite consensus sequences and their predicted frequency in satellite arrays based on consensus sequences. **D)** Histograms plotting the number of various A/T run lengths per 10 kb bin along portions of additional genomic contigs containing either *M. musculus* major satellite or *M. pahari* pi satellite arrays, indicated by tandem arrows above plots. **E)** The average number per 10 kb bin of all possible 4W and 5W and all present 6W sequences within the major and pi satellites arrays shown in Figure S2D and Figure 2E. **F)** Alignment of the 2EZD structure (magenta) and the corresponding AlphaFold3 prediction (green) with nucleotides noted along the DNA backbone. **G)** Summary of results of the *in silico* competitive binding assay, including when A/T sequence orders are reversed along the dsDNA. Includes results from Figure 2G. **H)** Part of an AlphaFold3 structure prediction highlighting how the AT-hook binds the minor groove of the sequence AAAAAT. The AT-hook (magenta) spans from A2 to T6 within the A/T-run. Model 0 of 5' GGGGAAAAATGGGGATATTAGGGG 3'. **I)** Histograms plotting the number of various $A_nT$ sequences per 10 kb bin along the same genomic contigs as in panel D.

**Table S1. Summary of *in silico* competitive binding assays.** Each row represents one of the five models ("Model output", 0-4) outputted by AlphaFold3 for a single competitive binding assay. For each model, the input DNA and Cbx2 AT-hook peptide sequence are listed. The column "Competing A/T runs" lists the two A/T runs in the input DNA sequence, and the column "AT-hook bound A/T run" lists which of these two A/T runs bound the AT-hook. Confidence scores for each model are also listed (ipTM and pTM). The final column lists the four to five nucleotide sequences that the AT-hook spans for AlphaFold models containing AAAAAT. n.d. = not determined, (A?) or (T?) = uncertain whether the A/T hook spans this nucleotide.

Figure S1

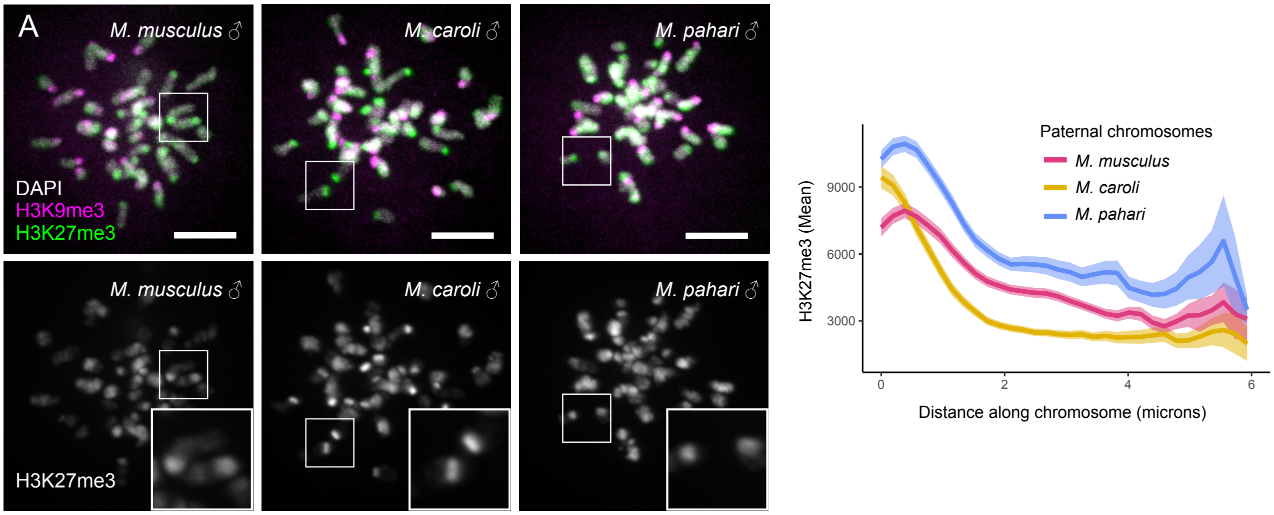
A
M. musculus ♂
M. caroli ♂
M. pahari ♂
DAPI
H3K9me3
H3K27me3
M. musculus ♂
M. caroli ♂
M. pahari ♂
H3K27me3
Paternal chromosomes
M. musculus
M. caroli
M. pahari
H3K27me3 (Mean)
9000
6000
3000
0
2
4
6
Distance along chromosome (microns)

### Figure S2

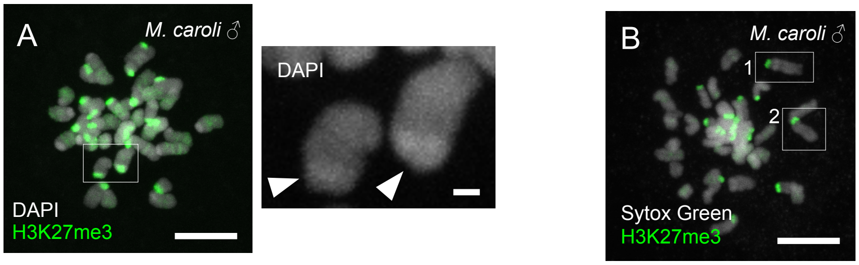


C

major satellite consensus

GAG**AAAT**GCACACTGAAGGACCTGG**AATAT**GGCGAG**AAAA**CTG**AAAAT**CACGG**AAAAT**GAG**A**
**AATA**CACAC**TTTA**GGACGTG**AAATAT**GGCGAGG**AAAA**CTG**AAAAA**GGTGG**AAAATTT**AG**AAA**
**T**GTCCACTGTAGGACGTGG**AATAT**GGCAAG**AAAA**CTG**AAAAT**CATGG**AAAAT**GAGAAACATC
CACTTGACGACTTG**AAAAAT**GACG**AAAT**CAC**TAAAAAA**CGTG**AAAAAT**

pi satellite consensus

CATGATTCACTCTG**TTTTT**CATGAG**TTTT**GTGTG**TAAAA**CAAGTG**AATTT**C**TTAAA**GATC**TA**
**TTA**GACACATTGTAGAG**ATTTTT**GTAGAACAAGC**ATAT**G**AATAT**GAG**TTTA**GTTCC**TAAAAT**
**A**CTGG**TTATT**CTATG**AAAA**CATTCCAC**AATT**CTTGTTCAGAGC**AATAA**GTACAACTATCTGC
ATT

| A/T run | Number per Major satellite consensus sequence | Predicted number per 10kb in Major satellite array | Number per Pi satellite consensus sequence | Predicted number per 10kb in Pi satellite array |
|---|---|---|---|---|
| 4W | 7 | 299 | 5 | 265 |
| 5W | 8 | 342 | 8 | 423 |
| 6W | 3 | 128 | 1 | 53 |
| 7W | 2 | 85 | 1 | 53 |
| 8W | 0 | 0 | 0 | 0 |
| 9W | 0 | 0 | 0 | 0 |
| ≥10W | 0 | 0 | 0 | 0 |
| Total | 20 | 855 | 15 | 794 |

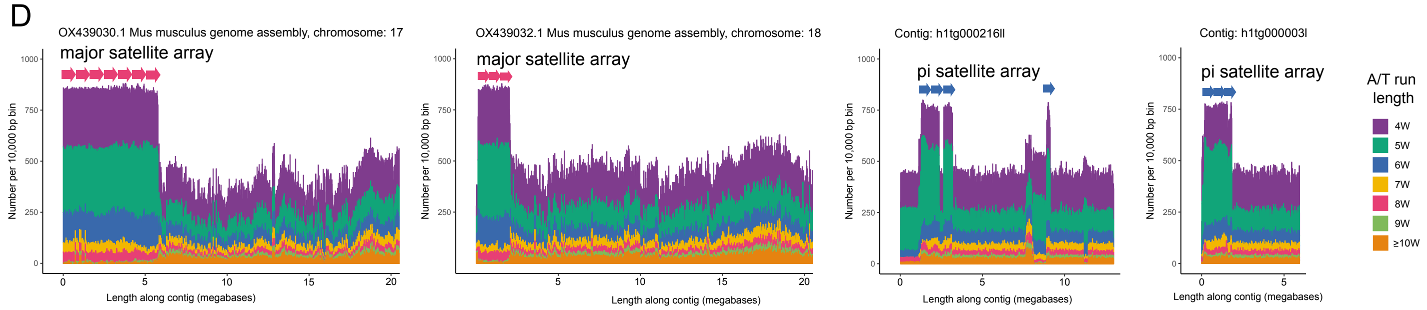


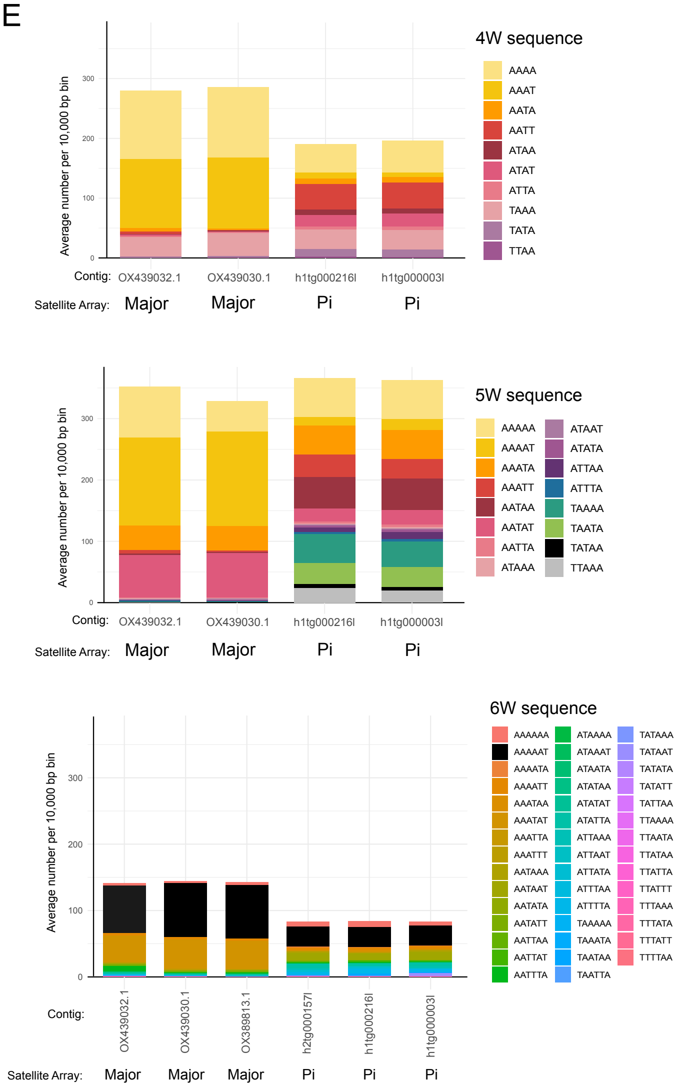


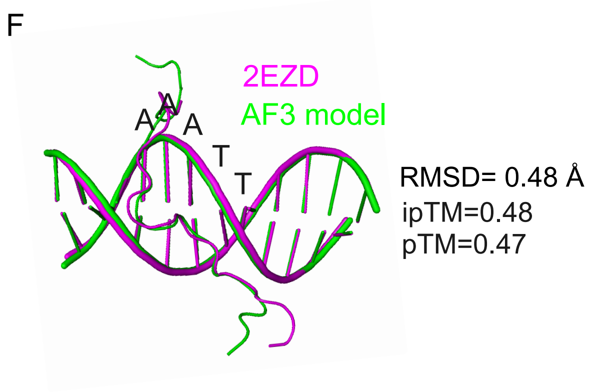


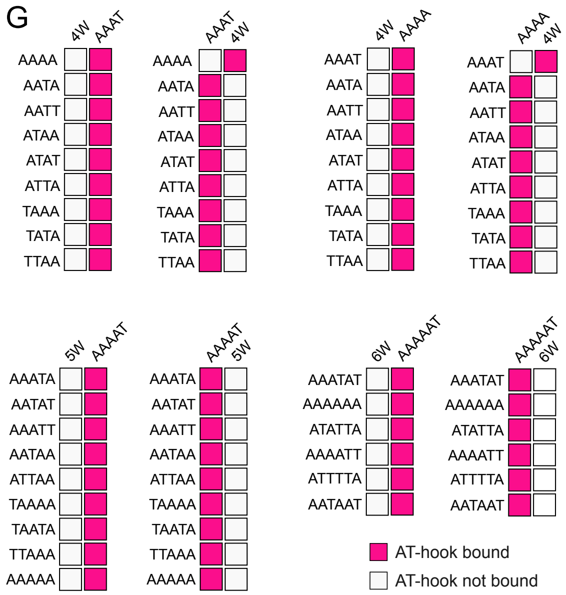

H

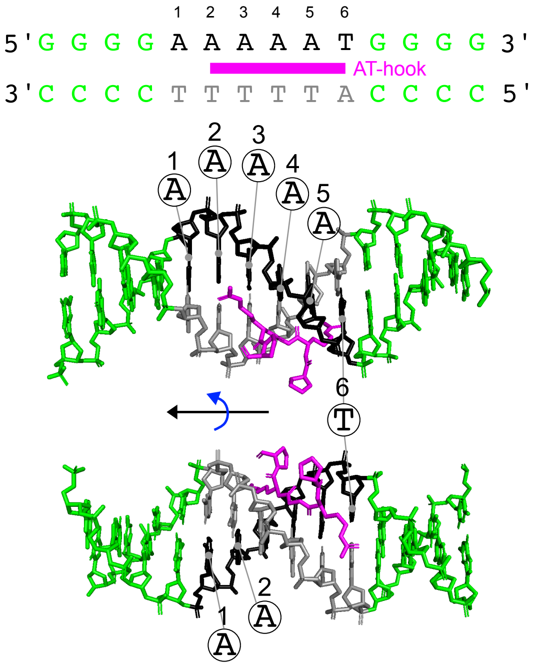
1 2 3 4 5 6
5' G G G G A A A A A T G G G G 3'
AT-hook
3' C C C C T T T T T A C C C C 5'
1 A
2 A
3 A
4 A
5 A
6 T
1 A
2 A

I

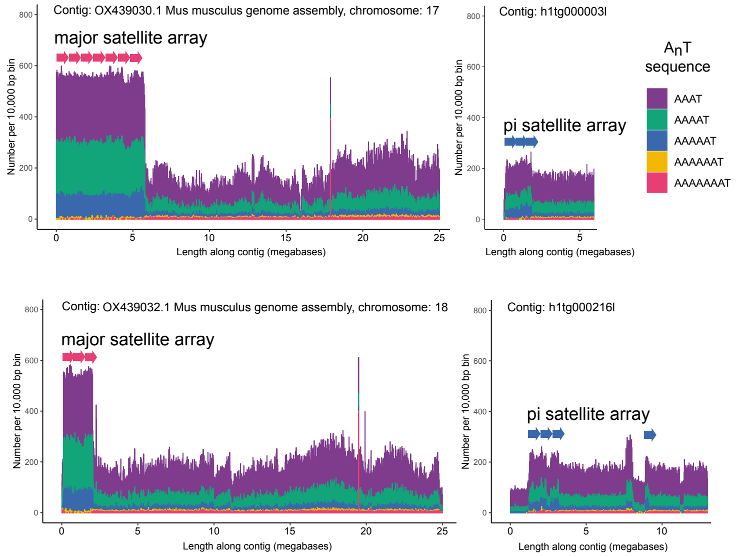
Contig: OX439030.1 Mus musculus genome assembly, chromosome: 17
major satellite array
Contig: h1tg000003l
pi satellite array
Contig: OX439032.1 Mus musculus genome assembly, chromosome: 18
major satellite array
Contig: h1tg000216l
pi satellite array
Number per 10,000 bp bin
Length along contig (megabases)
$A_nT$ sequence
AAAT
AAAAT
AAAAAT
AAAAAAT
AAAAAAAT
